# Electro-acupuncture Alleviates METH Withdrawal-induced Spatial Memory Deficits by Restoring Astrocyte-drived Glutamate Uptake in dCA1

**DOI:** 10.1101/2020.05.20.106153

**Authors:** Pengbo Shi, Zhaosu Li, Xing Xu, Jiaxun Nie, Dekang Liu, Qinglong Cai, Yonghua Zhao, Yun Guan, Feifei Ge, Xiaowei Guan

**Author notes:** Shi P, Li Z, Xu X and Nie J contributed equally to this work. Correspondence: Xiaowei Guan, MD, PhD, Department of Human Anatomy and Histoembryology, Nanjing University of Chinese Medicine, 138 Xianlin Ave, Nanjing 210023, China. Correspondence: Feifei Ge, PhD, Department of Human Anatomy and Histoembryology, Nanjing University of Chinese Medicine, 138 Xianlin Ave, Nanjing 210023, China.

## Abstract

Methamphetamine (METH) is frequently abused drug and produces cognitive deficits. METH could induce hyper-glutamatergic state in the brain, which could partially explain METH-related cognitive deficits, but the synaptic etiology remains incompletely understood. To address this issue, we explored the role of dCA1 tripartite synapses and the potential therapeutic effects of electro-acupuncture (EA) in the development of METH withdrawal-induced spatial memory deficits in mice. We found that METH withdrawal weakened astrocytic capacity of glutamate (Glu) uptake, but failed to change Glu release from dCA3, which lead to hyper-glutamatergic excitotoxicity at dCA1 tripartite synapses. By restoring the astrocytic capacity of Glu uptake, EA treatments suppressed the hyper-glutamatergic state and normalized the excitability of postsynaptic neuron in dCA1, finally alleviated spatial memory deficits in METH withdrawal mice. These findings indicate that astrocyte at tripartite synapses might be a key target for developing therapeutic interventions against METH-associated cognitive disorders, and EA represent a promising non-invasive therapeutic strategy for the management of drugs-caused neurotoxicity.

## INTRODUCTION

Methamphetamine (METH) is a frequently abused drug and often produces cognitive deficits, but there is lack of therapeutic approach to treat METH addiction and related cognitive deficits partially due to the less understanding of its etiology. It has been believed that METH could elicit endogenous glutamate (Glu) ^1^, which could partially explain METH-related addiction vulnerability and cognitive deficits ^2, 3^. Managing the homeostasis of extracellular Glu levels may protect the brain from excitotoxicity by drugs such as METH ^4^. However, the synaptic mechanism underlying METH-associated hyper-glutamatergic excitotoxicity remain unclear.

In the brain, astrocyte enwraps the neuronal components of synapse, which forming “tripartite synapse”. The hippocampus, specifically the dorsal CA1 (dCA1), plays key roles in cognitive function ^5, 6^ and METH neurotoxicity ^7, 8^. In the dCA1, tripartite synapse is constructed by the terminal of glutamatergic pyramidal neurons from dCA3, the spine of pyramidal neurons in the dCA1, and astrocytic processes. At tripartite synapse, astrocytes modulate Glu homeostasis by taking up the Glu at the synaptic cleft ^9, 10^. Based on the reactive statuses, astrocytes are divided into A1-like and A2-like astrocytes ^11^, which exhibit different physiological and pathological functions in the brain. Generally, A1-like astrocytes lost most normal astrocytic functions, but gained new neurotoxic function ^12^. There exists a perfect system for the uptake and metabolism of the Glu in the astrocytes at the tripartite synapse, such as glutamate transporter 1 (GLT-1), glutamate/aspartate-transporter (GLAST) and glutamine synthase (GS). GLT-1 and GLAST in the astrocytes are responsible for transmitting the Glu from cleft to the astrocytic plasm, while GS converts Glu into Glutamine (Gln) before being transported to neurons ^13, 14^.

Traditional Chinese medicine consider that “Governor Vessel (GV)” converged “Yang channels” over the whole body into the brain. Accordingly, GV stimulation may exert beneficial effects in brain diseases. Electro-acupuncture (EA) has been used to treat brain disorders, including cognitive dysfunctions ^15, 16^. In clinic, EA at the acupoints of Yintang (GV29) and Baihui (GV20) were used to treat depression ^17^, Alzheimer’s disease ^18^ and Tourette syndrome ^19^. Then, we wondered whether and how EA at GV29 and GV20 influence METH-related cognitive deficits.

In the present study, we hypothesize that METH withdrawal impairs the astrocytes-dependent

Glu uptake at tripartite synapses in the dCA1, which contribute to the spatial memory deficits in mice. EA at acupoints of Yintang and Baihui rescue the spatial memory deficits in METH withdrawal mice, partially by restoring the astrocytic capacity of Glu uptake.

## RESULTS

### METH withdrawal caused hyper-glutamatergic status in dCA1 and spatial memory deficits in mice, and EA treatments ameliorated these damages

METH withdrawal mice (METH mice) and saline control mice (SAL mice) were established as Figure 1 A1. During METH withdrawal period, some mice were subjected to EA treatments (M + EA mice) at the acupoints of GV29 and GV20 or Sham EA (SEA) treatments at “placebo-point” in mice (M + SEA mice, Figure 1 A2). Y-maze, Morris water maze (MWM) and object location memory (OLM) assays were used to evaluate short-term spatial, long-term spatial and location memory, respectively. In Y-maze test, METH mice exhibited less preference to novel arm (NA) of maze than that of SAL mice (p = 0.0153, n = 22, Figure 1 B1), while EA treatments increased the preference to NA in METH mice (p = 0.0063, n = 22, Figure 1 B2) when compared with M + SEA mice. In the MWM test, METH mice spent more time to find the platform in the escape latency test (T1, p = 0.0024, n = 20, Figure 1 C1), but less time to explore in the quadrant of hidden platform in the probe test (T2, p = 0.0016, n = 20, Figure 1 C2) when compared to SAL mice. There are no differences in escape latency between METH and SAL mice during the training period (Day 1-3, p > 0.05, n = 20, Figure 1 C1). EA treatments did not change the escape latency in the T1 (p = 0.4883, n = 20, Figure 1 C3), but increased the exploring time in the target quadrant of T2 (p = 0.0044, n = 20, Figure 1 C4) when compared to M + SEA mice. In the OLM test (Supplemental Figure 1 A), METH withdrawal decreased the discrimination index (p = 0.0301 vs. SAL, n = 22, Supplemental Figure 1 B1), and EA treatments failed to reverse the impaired location memory in METH mice (p = 0.3092 vs. M + SEA, n = 20, Supplemental Figure 1 B2). Both METH withdrawal (p = 0.3114 vs. SAL, n = 22, Supplemental Figure 1 C1) and EA treatments (p = 0.7919 vs. M + SEA, n = 20, Supplemental Figure 1 C2) did not alter the total distance travelled by mice in the test box.

**Figure 1.**
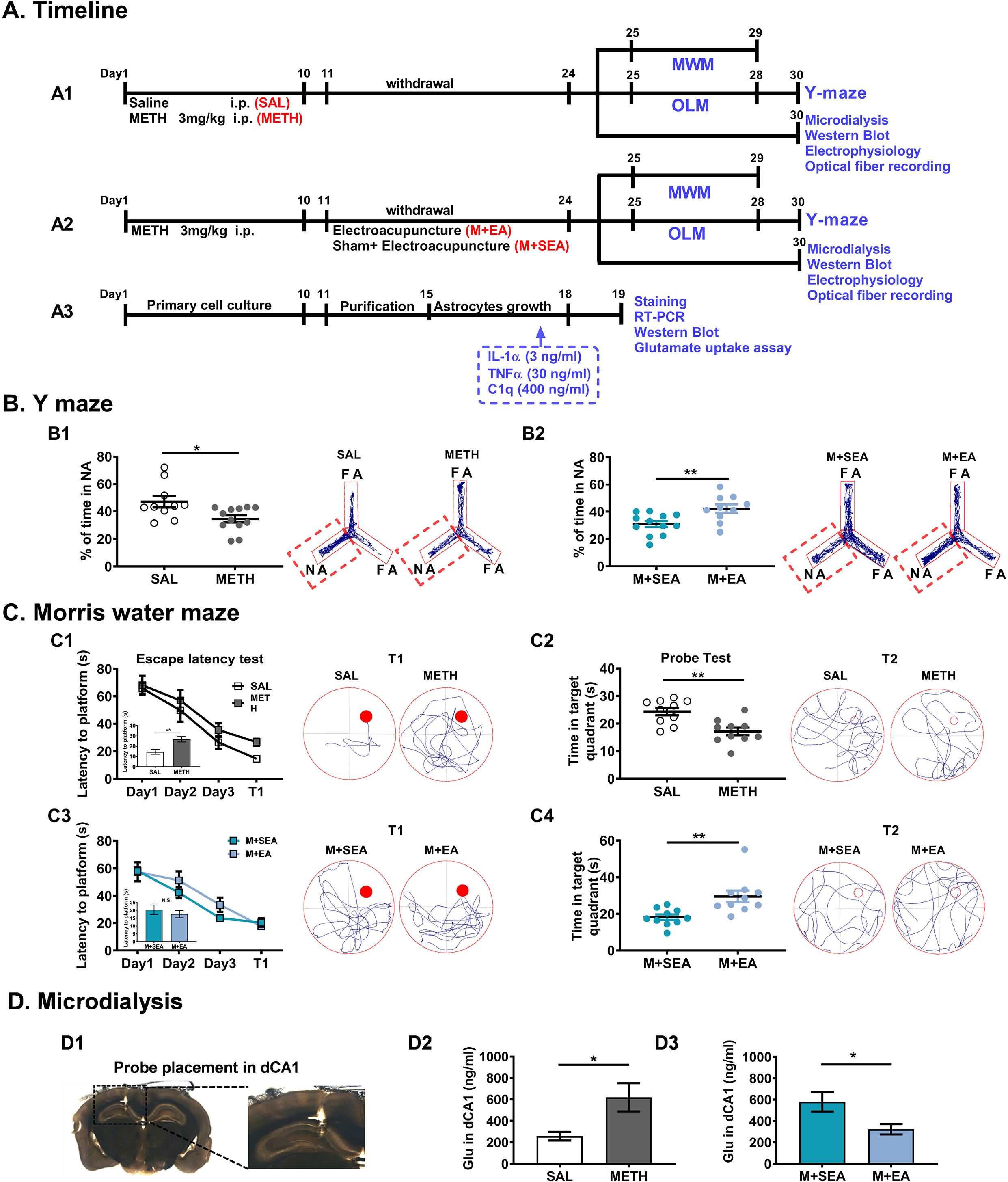
EA at Yintang (GV29) and Baihui (GV20) alleviated METH withdrawal-induced spatial memory deficits in mice and hyper-glutamatergic state in dCA1. A, Outline of the experimental protocol. Experimental design and timeline for *in vivo* (A1-2) and *in vitro* (A3) studies. **B**, Y maze test. (B1), Time (% total) spent in the novel arm (NA) (left) and representative traces (right) of SAL and METH mice during the Y maze test. (B2), Time (%) spent in the NA and representative traces (right) during the Y maze test of M+SEA and M+EA mice. **C**, Morris water maze. (C1, C3), Learning curves in hidden platform training (Day 1-3) and latency to the platform during the escape latency test (T1, left); Representative swimming paths during T1 test (right). (C2, C4), Target quadrant time (left) and representative swimming paths (right) in the MWM probe test. **D**, Microdialysis study. (D1), Example images of location of microdialysis probe in the dCA1. (D2), Glu levels in the dCA1 of SAL and METH mice. (D3), Glu levels in the dCA1 of M+SEA and M+EA mice. Data are Mean ± S.E.M.

To detect the extracellular Glu and Gln concentrations at the dCA1, *in vivo* microdialysis and liquid chromatography-mass spectrometry (LC-MS) were used here (Figure 1 D1). When compared to SAL mice, Glu levels were increased at the dCA1 in METH mice (p = 0.0187, n = 11 Figure 1 D2), while EA treatments suppressed the extracellular Glu levels in METH mice (p = 0.043 vs M + SEA mice, n = 11, Figure 1 D3). In parallel, extracellular Gln levels at the dCA1 were not influenced by either METH withdrawal (p = 0.6327, n = 12, Supplemental Figure 2 A) or EA treatments (p = 0.1142, n = 13, Supplemental Figure 2 B), when compared with corresponding control mice.

### METH withdrawal weakened astrocytic capacity of Glu uptake at dCA1 tripartite synapses, and EA treatments normalizing the function of astrocyte

Complement component 3 (C3), S100A10 and glial fibrillary acidic protein (GFAP) were used as markers for the A1-like, A2-like and all astrocytes respectively here. As previously reported, reactive astrocyte shows thicker cellular processes and larger body compared to the resting ones ^20^. When compared to SAL mice, METH withdrawal significantly increased the proportion of C3-positive A1-like astrocytes (p = 0.0016, n = 12) and reactive astrocytes (p = 0.0007, n = 12), but failed to alter the total number of GFAP-positive astrocyte (p = 0.0961, n = 12, Figure 2 A1), the proportion of S100A10-positive A2-like astrocytes (p = 0.1921, n = 12, Supplemental Figure 3 A1) and the protein levels of S100A10 (p = 0.9050, n = 6, Supplemental Figure 3 B1) at dCA1. In parallel, the protein levels of C3 (p = 0.0353, n = 6) and GFAP (p = 0.0155, n = 6) were increased at dCA1 in METH mice, as compared to SAL mice (Figure 2 A2). Compared with SEA treatments, EA treatments attenuated the proportion of C3-positive A1-like astrocytes (p = 0.0015, n = 12) and reactive astrocytes (p = 0.0278, n = 12, Figure 2 A3), as well as the protein levels of C3 (p = 0.0027, n = 6) and GFAP (p = 0.0054, n = 6, Figure 2 A4) in the dCA1 of METH mice, but failed to alter either the proportion of S100A10-positive A2-like astrocytes (p = 0.1985, n = 12, Supplemental Figure 3 A2) or the protein levels of S100A10 (p = 0.4091, n = 12, Supplemental Figure 3 B2) at dCA1.

**Figure 2.**
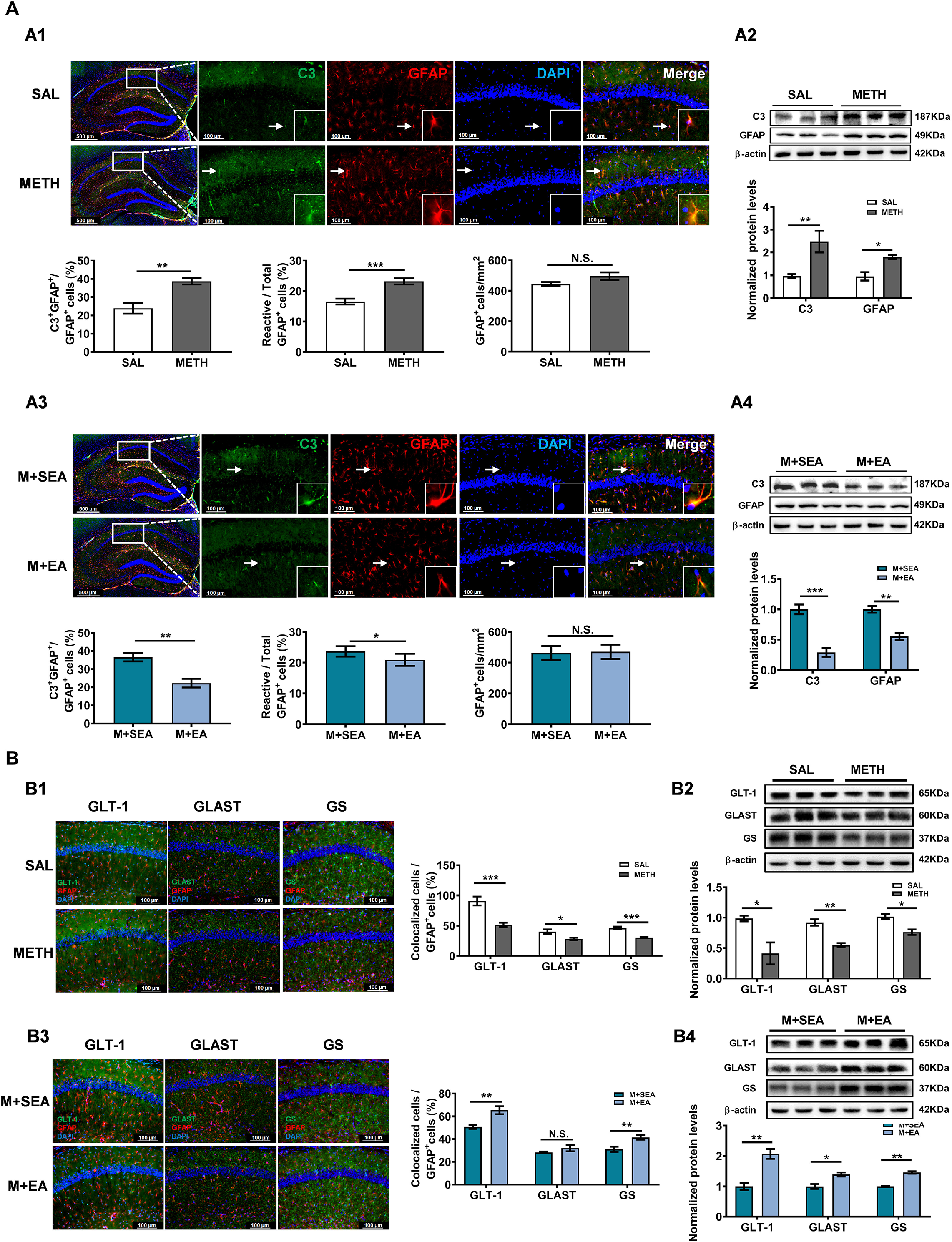
EA rescued METH-related neurotoxicity via normalizing the function of astrocyte. A. Expression of C3 in the dCA1 during METH withdrawal. (A1), Top panels: immunohistochemistry staining for C3 (green), GFAP (red) and DAPI (blue) in the hippocampus of SAL and METH group mice. White arrows indicate representative cells which were showed at a higher power view in the white box. Bottom panels: left, the proportion of C3 and GFAP double-label cells; middle, the proportion of reactive astrocytes; right, numbers of GFAP-positive cells per mm^2^ in SAL and METH groups. (A2), Western blots (top) and quantification (bottom) of C3 and GFAP protein levels in the dCA1 of SAL and METH groups. (A3), Top panels: immunohistochemistry staining for C3 (green), GFAP (red) and DAPI (blue) in the hippocampus of M+SEA and M+EA group mice. Bottom panels: left, the proportion of C3 and GFAP double-label cells; middle, the proportion of reactive astrocytes; right, number of GFAP-positive cells per mm^2^. (A4), Western blots (top) and quantification (bottom) of C3 and GFAP protein levels in the dCA1 of M+SEA and M+EA group. **B,** GLT-1, GLAST and GS expression levels in the dCA1 during METH withdrawal. (B1), Immunohistochemistry staining (left, GLT-1/GLAST/GS-green; GFAP-red and DAPI-blue) and colocalized ratio (right) for GLT-1, GLAST and GS in the hippocampus of SAL and METH group mice. (B2), Western blots (top) and quantification (bottom) of GLT-1, GLAST and GS protein levels in the dCA1 of SAL and METH group. (B3), Immunohistochemistry staining (left) and colocalized ratio (right) for GLT-1, GLAST and GS of M+SEA and M+EA group. (B4), Western blots (top) and quantification (bottom) of GLT-1, GLAST and GS protein levels in the dCA1 of M+SEA and M+EA group. Data are Mean ± S.E.M.

The molecules involved in uptake and metabolism of Glu in the astrocytes were detected here. As shown in Supplemental Figure 4 A, most of GS in the dCA1 was co-expressed with GFAP-positive astrocytes, but few with NeuN-marked neurons (p = 0.0002, n = 6). METH withdrawal significantly decreased the percentage of GLT-1 (p = 0.0007, n = 12), GLAST (p = 0.0204, n = 12) and GS-positive astrocytes (p = 0.0001, n = 12, Figure 2 B1), as well as the protein levels of GLT-1 (p = 0.0371, n = 6), GLAST (p = 0.0037, n = 6) and GS (p = 0.0138, n = 6, Figure 2 B2), when compared with that of SAL mice. Compared with SEA treatments, EA treatments enhanced the percentage of GLT-1 (p = 0.0033, n = 12) and GS (p = 0.0074, n = 12), but failed to change that of GLAST (p = 0.2087, n = 12, Figure 2 B3) astrocytes in METH mice. In parallel, EA treatments also increased the protein levels of GLT-1 (p = 0.0060, n = 6), GLAST (p = 0.0191, n = 6) and GS (p = 0.0004, n = 6) at dCA1 in METH mice (Figure 2 B4).

As shown in Figure 3 A1, 96.22% purified astrocytes were obtained in the primary cultured cells (p < 0.0001, n = 12) from mice cortex. To assess the capacity of A1-like astrocytes in the process of Glu uptake, A1-like phenotype astrocyte were induced by incubation with interleukin 1α (Il-1α), tumor necrosis factor (TNF), complement component 1 and subcomponent q (C1q) ^12^. After 24-h induction, astrocytes showed a stable A1-like phenotype, as indicated by the 78.53% of C3-positive cells (p = 0.0012, n = 6) and increased C3 mRNA levels (p < 0.0001, n = 6, Figure 3 A2) when compared with controls (Cont astrocyte). *In vitro*, A1-like astrocytes cleared much less Glu in the culture medium compared to the Cont astrocytes (Figure 3 A3, F _(1,4)_ = 25.02, p = 0.0075, two-way ANOVA). In parallel, the percentage of GLT-1-positive value (p = 0.0056, n = 20) and GS-positive value (p = 0.0351, n = 20, Figure 3B1), the protein levels of GLT-1 (p = 0.0145, n = 6, Figure 3 B2) and GS (p = 0.0327, n = 6 Figure 3B2) were significantly decreased in cultured A1-like astrocytes. While, no deference in GLAST-positive value (p = 0.2223, n = 20, Figure 3 B1) and protein levels of GLAST (p = 0.1918, n = 20, Figure 3 B2) were observed in cells between Cont and A1-like astrocytes.

**Figure 3.**
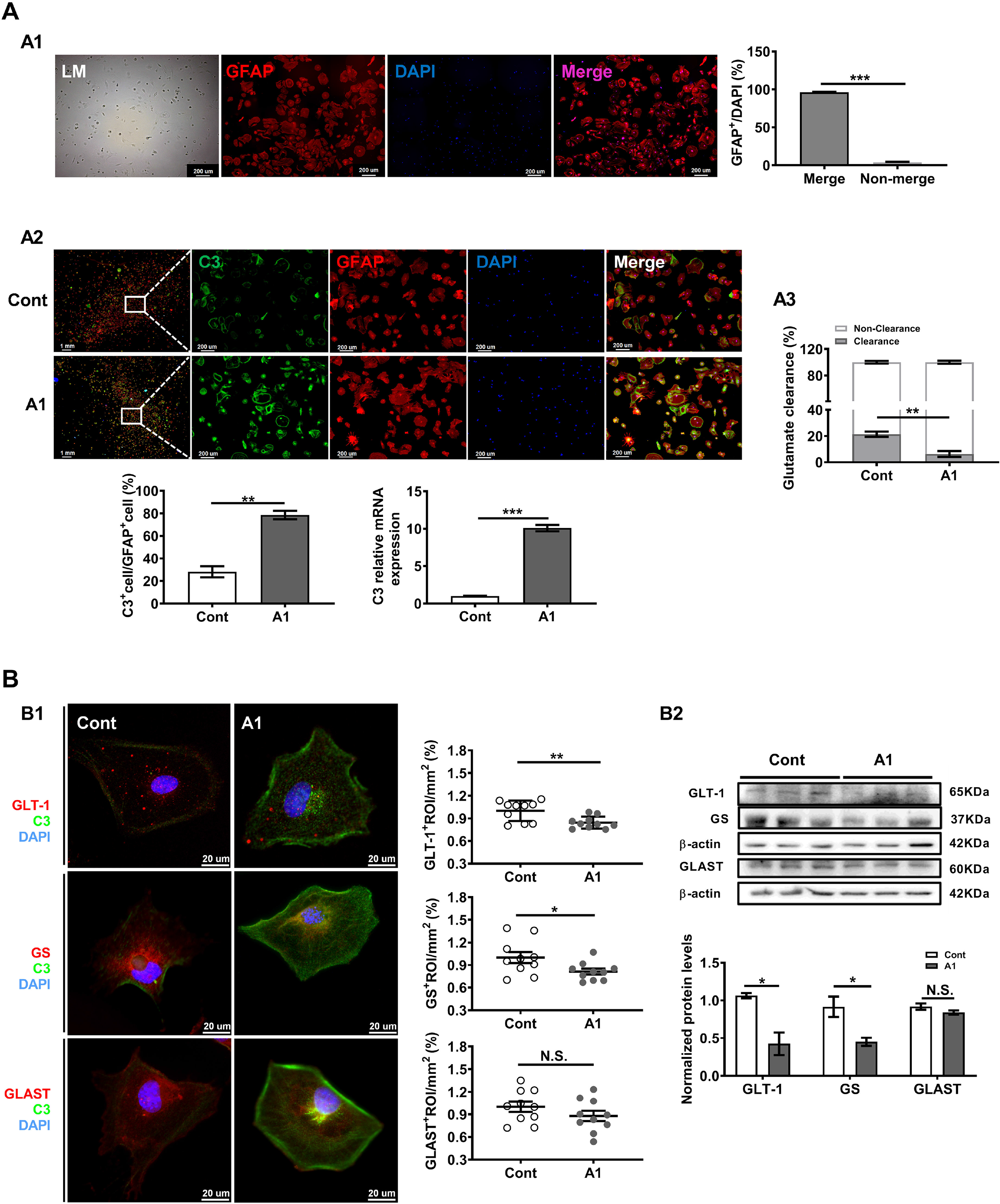
A1-like astrocytes showed decreased capacity of Glu uptake. A, Glu clearance in primary cultured astrocytes. (A1), Representative micrographs (left) and quantification (right) for GFAP staining in primary astrocytes. (A2), Top panels: immunohistochemistry micrographs for C3 (green), GFAP (red) and DAPI (blue) of control and A1 group. Bottom panels: left, ratio of C3 positive astrocytes; right, mRNA expression levels of C3. (A3), Ratio of Glu clearance after 1h application of Glu (100μM) in the control and A1 group. **B,** GLT-1, GLAST and GS expression in the primary cultured astrocytes. (B1), Immunohistochemistry micrographs (left, GLT-1/GLAST/GS-red; C3-green and DAPI-blue) and quantification (right) for GLT-1, GS and GLAST of control and A1 group. (B2), Western blots (top) and quantification (bottom) of GLT-1, GS and GLAST protein levels of control and A1 group. Data are Mean ± S.E.M.

### METH withdrawal and EA treatment failed to influence presynaptic Glu transmission from dCA3

To examine the presynaptic Glu transmission at dCA1 tripartite synapses, the activities of presynaptic neurons were recorded in the Schaffer projections from dCA3 to dCA1. Here, c-Fos was used as marker for assessing the neuron activities, and vesicular glutamate transporter (VGLUT) was used as marker for evaluating the capacity of Glu release from presynaptic neurons, and synapsin 1 (SYP1) was used as a marker to show the terminals of presynaptic neurons. As shown in the Figure 4 A, neither METH withdrawal (p = 0.8054 vs. SAL mice, n = 12) nor EA treatments (p = 0.6139 vs M + SEA mice, n = 12) altered c-Fos staining at dCA3 when compared to that of corresponding controls. In parallel, both METH withdrawal (p =0.3052 vs. SAL mice, n = 12) and EA treatments (p = 0.2459 vs. M + SEA mice, n = 12, Figure 4 B) failed to change VGLUT-1 staining in terminals of neurons from dCA3 in the dCA1.

**Figure 4.**
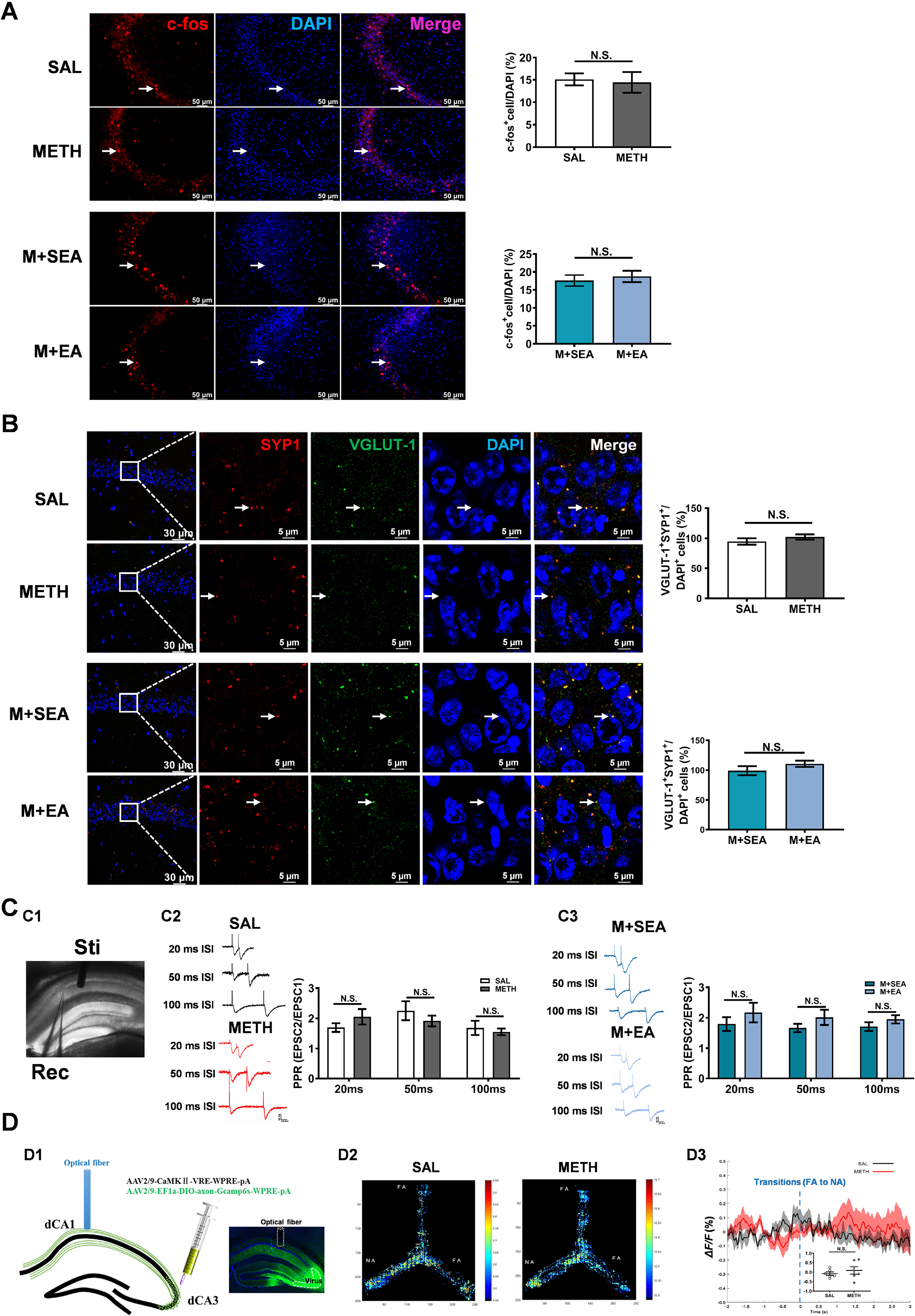
METH and EA treatments did not affect presynaptic glutamatergic transmission from dCA3. A, Expression of c-Fos in the dCA3 after exposure to behavior test for 90 min. Left, representative images of c-Fos (red) and DAPI (blue) labeling in the dCA3. Right, the numbers of c-Fos positive cells per mm^2^. White arrows indicate positive staining of c-Fos. **B,** Immunohistochemistry micrographs (left, SYP1-red; VGLUT-1-green and DAPI-blue) and colocalization (right) of SYP1 and VGLUT-1. White arrows indicate colocalizations. **C,** Paired-pulse ratio (PPR) in post-synaptic dCA1 neurons. (C1), Position of stimulatory electrode and recording pipette. (C2), Representative traces (left) and PPR (right) in the dCA1 neurons evoked by test stimulation with interpulse intervals of 20, 50 and 100 ms in SAL and METH group mice. (C3), Representative traces (left) and PPR (right) in the dCA1 neurons evoked by test stimulation with interpulse intervals of 20, 50 and 100 ms in M+SEA and M+EA group. **D**, Fiber photometry used to measure dCA3 axonal calcium signals in the dCA1 during Y maze test in free-moving mice. (D1), Left, diagram shows experimental setup; Right, representative images of virus expression. (D2), GCaMP fluorescence intensity with respect to animal’s position during Y maze test of SAL and METH group mice. (D3), Averaged fluorescence intensity at the points of transitions between FA to NA of SAL and METH group mice; Inset: scatter plot shows averaged area under the curve after entering into NA. Data are Mean ± S.E.M.

To assess the electrophysiological characteristics of presynaptic neurons in the dCA3, paired-pulse ratio (PPR) was recorded in post-synaptic dCA1 neurons (Figure 4 C1). There was no difference in PPR between METH and SAL mice (F _(2, 60)_ = 1.466, p = 0.2390, two-way ANOVA, Figure 4 C2), and between M + SEA and M + EA mice (F _(2, 63)_ = 0.058, p = 0.9437, two-way ANOVA, Figure 4 C3).

To examine the glutamatergic transmission from dCA3 to dCA1 *in vivo*, a mixture of viruses (AAV2/9-CaMKⅡ-VRE-WPRE-pA and AAV2/9-EF1a-DIO-axon-Gcamp6s-WPRE-pA) were injected into the dCA3. The injected viruses specifically transfected glutamatergic pyramidal neurons in the dCA3, and the GCaMP6 could be transported from the dCA3 to the dCA1 along the axon of Schaffer fibers (Figure 4 D1). During Y maze test, the real-time calcium signals of Schaffer fibers terminals were recorded in the dCA1. In free-moving mice of Y-maze, no differences in the real-time pre-synaptic calcium signals between METH and SAL mice were recorded in the dCA1 (Figure 4 D2). In parallel, the ratio of the fluorescence intensity to the resting fluorescence intensity (ΔF/F) even were not affected by METH withdrawal in mice (p = 0.4569 vs. SAL mice, n = 12, Figure 4 D3).

### METH withdrawal triggered postsynaptic neurons at dCA1 tripartite synapses, and EA treatments suppressed the Glu excitotoxicity

METH withdrawal enhanced the percentage of c-Fos-positive pyramidal neurons at dCA1 when compared to that of SAL mice (p = 0.0396, n =12, Figure 5 A1), while EA treatment reversed the changes in c-Fos of dCA1 (p = 0.0491, n =12, Figure 5 A2). In addition, increases in amplitude of miniature excitatory postsynaptic currents (mEPSCs), but not the frequency (p = 0.6905 vs. SAL mice, n = 26) were recorded in the dCA1 of METH mice (p = 0.0103 vs. SAL mice, n = 26, Figure 5 B1). EA treatments attenuated the amplitude of mEPSCs in postsynaptic dCA1 neurons (p = 0.0486, n = 21) and has no influence on that of frequency (p = 0.3703, n =21, Figure 5 B2) in the dCA1 of METH mice.

**Figure 5.**
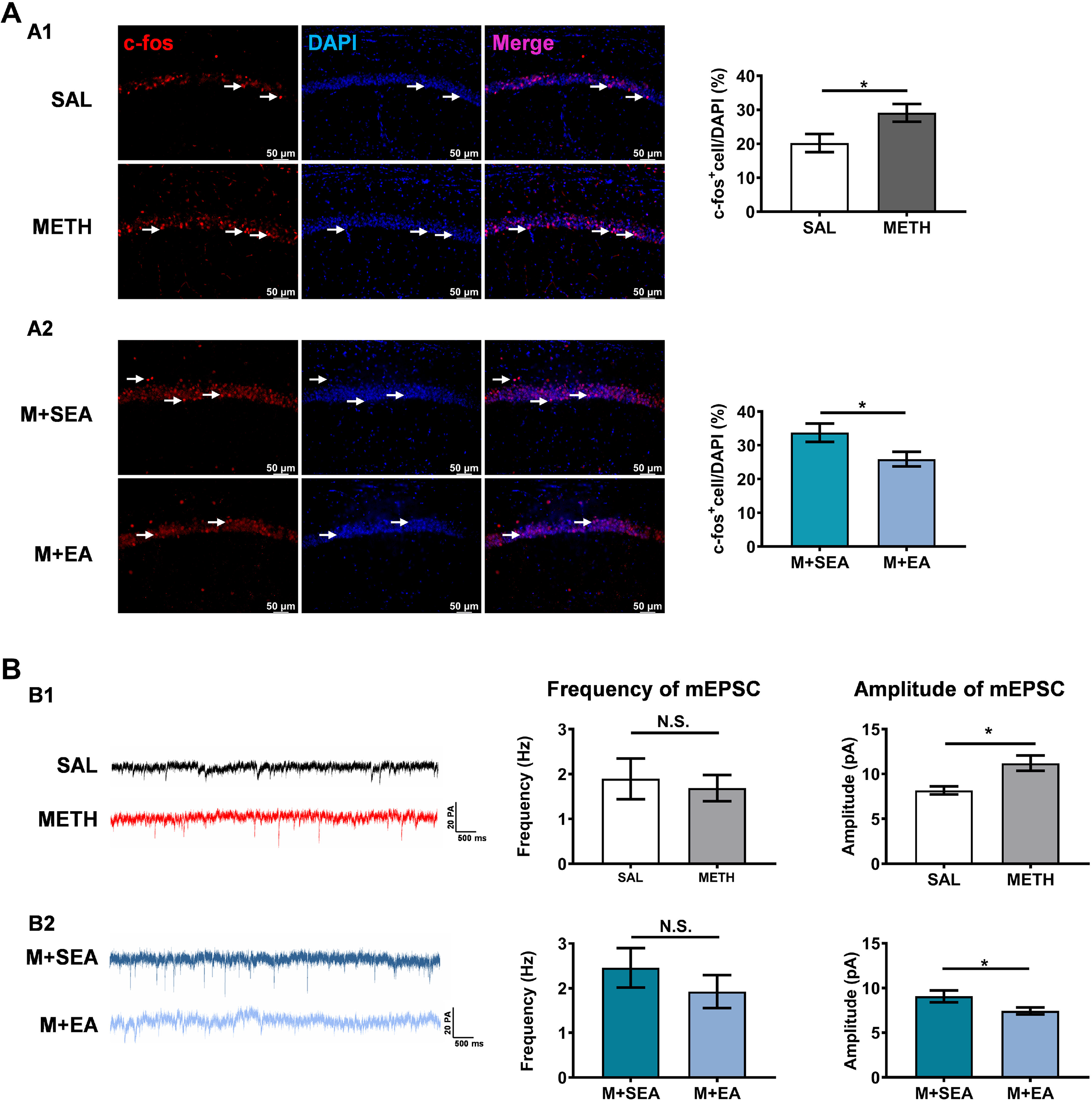
EA treatments alleviated METH withdrawal-induced glutamatergic excitotoxicity in dCA1. A, Induction of c-Fos expression in the dCA1 after exposure to behavior test for 90 min. Left, representative photomicrographs of c-Fos (red) and DAPI (blue) labeling in the dCA1. Right, the numbers of c-Fos positive cells per mm^2^ in each group. White arrows indicate positive staining of c-Fos. **B,** Representative traces and quantification of the frequency and amplitude of spontaneous miniature excitatory postsynaptic currents (mEPSCs) in CA1 neurons. Data are Mean ± S.E.M.

Taken together (Figure 6), our data demonstrated that ➀METH withdrawal reduce astrocytedependent Glu uptake at dCA1 tripartite synapses, ➁but did not alter glutamatergic transmission from CA3, ➂subsequently increase the Glu concentration at synaptic clefts of dCA1, ➃then lead to postsynaptic neuron hyperexcitability in the dCA1, which contribute to the spatial memory deficits. By restoring astrocytic capacity of Glu uptake, EA treatment at acupoints of Yintang (GV29) and Baihui (GV20) efficiently inhibited the hyper-glutamatergic state and normalized the activities of postsynaptic neuron at dCA1 tripartite synapse, then alleviated METH withdrawal-induced spatial memory deficits.

**Figure 6.**
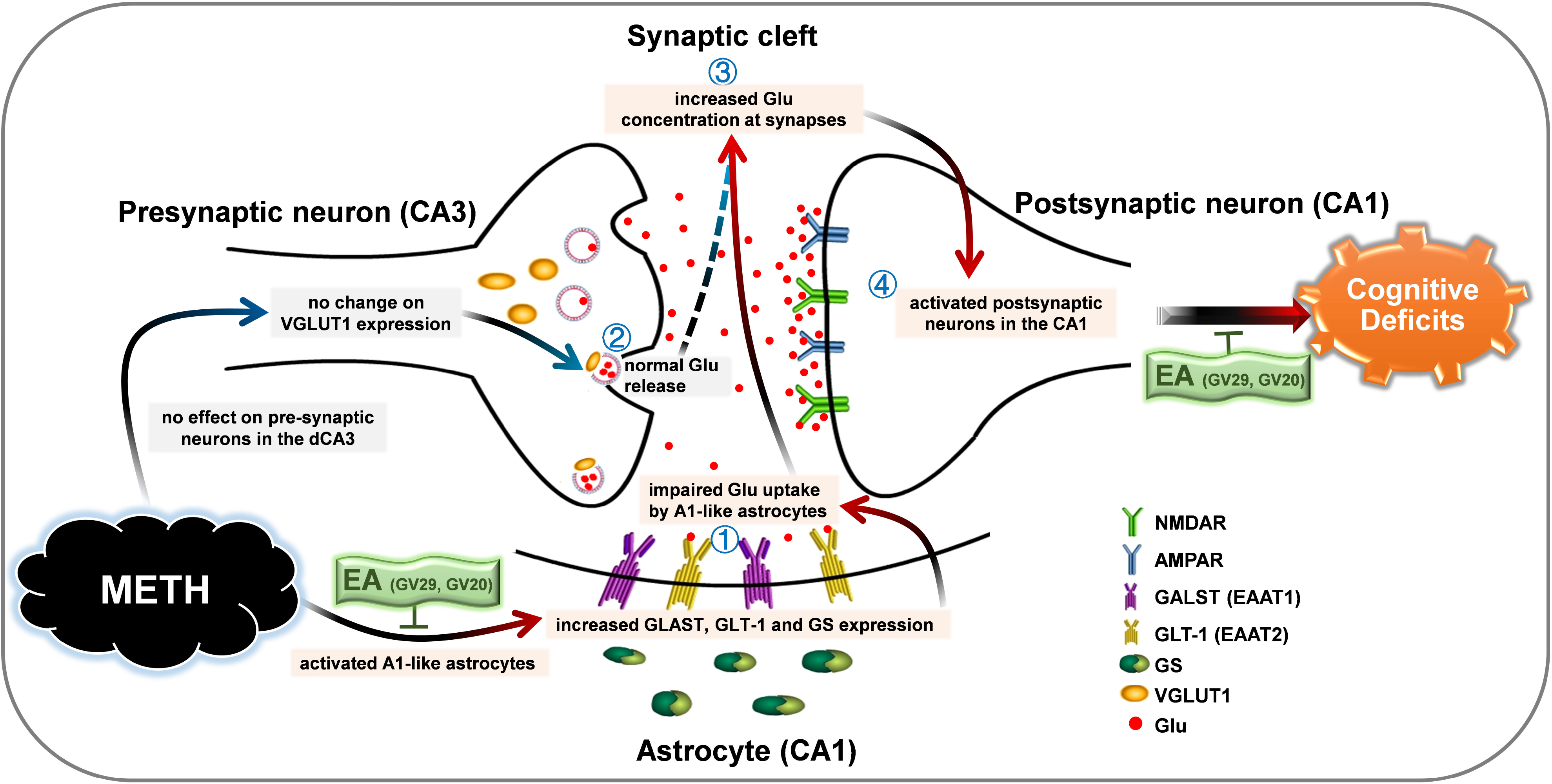
Schematic diagram of the present study. ➀ Glu uptake by astrocytes at the tripartite synapses of dCA1; ➁ Glu releasing from the presynaptic terminals of dCA3; ➂ Glu concentration at synaptic clefts in dCA1; ➃ Activities of postsynaptic neurons in dCA1.

## DISCUSSION

METH abusers often exhibit memory deficits, which triggers proclivity to addiction. Understanding of the etiology underlying METH-induced memory impairments is crucial for developing intervention approaches that improve cognitive performance and overall addiction. In this study, we found that METH withdrawal triggered Glu transmission at dCA1 tripartite synapse due to the reduction of astrocyte-dependent Glu uptake, which are associated with METH-impaired spatial memory in mice. By correcting the etiology of astrocyte, EA effectively ameliorate the spatial memory deficits in METH withdrawal mice.

### METH withdrawal causes a hyper-glutamatergic state at dCA1 synapses in mice

Glu is the primary excitatory neurotransmitter in the brain, and synaptic Glu homeostasis decide the efficiency of excitatory transmission in the brain. Excessive extracellular Glu levels, such as being found in METH abusers and METH-experienced animals, can lead to the overstimulation of postsynaptic neurons, which elicit excitotoxicity in the neurocircuitry and abnormal behavioral outcomes ^1, 9, 21–23^. In the current study, METH withdrawal mice exhibit both long-term and shortterm impaired spatial memory. Memory for spatial recognition is primarily mediated by the hippocampus, especially dCA1. Consistent with the hyper-glutamatergic states found in striatum ^8^, prefrontal cortex ^24^ and nucleus accumbens ^1^ by METH use, we found that extracellular Glu concentrations in the dCA1 consistently kept at high levels, and dCA1 pyramidal neurons were strongly activated in METH withdrawal mice. To assess the relationship of dCA1 Glu hyperactivity and the spatial memory deficits in METH withdrawal mice, EA at acupoints of Yintang (GV29) and Baihui (GV20) were performed to METH mice during the whole withdrawal period. We found that EA significantly suppressed the hyper-glutamatergic state, normalized the activities of postsynaptic neuron in dCA1, and alleviated the spatial memory deficits in METH mice as well. The EA treatment outcome has potential significance to inform clinical assessment of “anti-glutamate” therapeutic strategies for attenuating METH-related abnormal behaviors.

### The reduction of astrocyte-derived Glu uptake contribute to the excitotoxicity at dCA1 tripartite synapses in METH withdrawal mice

Although some studies have reported the glutamate-associated excitotoxic lesions in METH abusers or animals, the synaptic mechanisms underlying METH-induced hyper-glutamatergic state remain unclear. In the dCA1, the postsynaptic pyramidal neurons receive the glutamatergic afferents from dCA3, and astrocyte enwraps the pre- and post-neuronal components of synapse, which forming “tripartite synapse” ^14, 25^. Basal extrasynaptic Glu tone is maintained largely by presynaptic neuronal Glu release and astrocyte-mediated uptake of Glu ^26, 27^. Astrocytes undergo a dynamic shift between two different reactive states, A1 phenotype and A2 phenotype astrocytes. A1-like astrocytes lose many normal astrocytic functions but gain some new neurotoxic function ^12^. By contrast, A2-like astrocytes upregulate many neurotrophic factors and therefore has been thought of protective phenotype astrocyte ^12^. In the current study, we found that the percentage of A1-like astrocyte but not that of A2-like astrocytes were increased in dCA1 of mice during METH withdrawal period. *In vitro* experiments found that A1-like astrocyte indeed lost the capacity to taking up the extracellular Glu. In parallel, METH withdrawal failed to affect the activities of presynaptic Glu transmission from dCA3. These results indicate it was the disruption in astrocyte-drived Glu uptake by METH withdrawal but not the Glu release from dCA3 that account for the hyper-glutamatergic state in the dCA1 of mice.

Astrocyte transmit the Glu from extracellular tissue to the astrocytic plasm through two predominant transporter isoforms ^28^, GLAST and GLT-1 in rodents (homologs of EAAT1 and EAAT2 in humans, respectively). In addition, GS in astrocytes is responsible for converting Glu into Gln before being transported to neurons ^13, 14^. In adults, the GLT-1 is more effective with a higher turnover rate than GLAST, accounting for more than 90% of total uptake in the brain ^29^. In the current study, we found that GLT-1 and GS but not GLAST were decreased in astrocyte after being induced into A1-like phenotype *in vitro*. In addition, METH withdrawal *in vivo* decreased expressions of both GLT-1 and GLAST in dCA1, which is consistent with previous findings about GLT-1 and GLAST in other drugs of models such as cocaine and heroin ^27, 30^. Taken together, these results indicate that the dysfunctions in astrocytic glutamate transporting system, especially GLT-1 might be key pathological molecules that contribute to the weakened capacity of astrocyte-drived Glu uptake by METH withdrawal.

### EA treatments efficiently alleviate the spatial memory deficits in METH withdrawal mice by restoring astrocytic capacity of Glu uptake at dCA1

At present, there are no safe and effective drugs for treating METH abuse and related behaviors deficits. Acupuncture is one non-invasion approach of traditional Chinese medicine that has been practiced in the treatment of brain disorders for a long time. In clinic, Five-Point Protocol, a form of auricular acupuncture that are developed by the National Acupuncture Detoxification Association (NADA), have been used to prevent substance abusers from neural toxification and addiction ^31^. In addition, acupuncture has been shown to effectively inhibit heroin’s neurotoxic effects ^32^, suggesting that acupuncture may benefit for METH-associated neurotoxicity. The acupoints of GV29 and GV20 are considered to be the optimized acupoint modules for mental illness ^17^, but the therapeutic mechanism remains unclear. EA at the acupoints of GV29 and GV20 show a better efficacy in improving the spatial learning and memory ability in patients of Alzheimer’s disease ^33^. Thus, we proposed that EA might be an ideal therapies for METH-impaired spatial memory behaviors. In this study, EA were performed on GV29 and GV20 acupoints for 30 min once a day for 14 days in METH withdrawal mice. we found that long-term EA performance obviously alleviated METH withdrawal-induced spatial memory deficit behaviors in mice. Then, we explore the potential mechanisms underlying the therapeutic effects of EA on the abnormal behaviors by METH withdrawal. We found that EA can “calm down” the METH withdrawal-caused excitotoxicity at dCA1, mainly through reversing the enhanced A1-like phenotype astrocytes and rescuing the capacity of astrocyte-mediated glutamate transporting at tripartite synapses. As to the transporter molecules, EA treatments only increased GLT-1but not GLAST transporter at dCA1.

In conclusion, our findings demonstrate that the weakened capacity of astrocyte-drived Glu uptake contribute to METH withdrawal-caused hyper-glutamatergic excitotoxicity in dCA1. Targeting the process of Glu transmission in dCA1, especially by restoring the astrocytic capacity of Glu uptake, EA treatments efficiently rescue the spatial memory deficits in METH withdrawal mice. These findings indicate that astrocyte at tripartite synapses might be a novel target for developing therapeutic interventions against METH-associated cognitive disorders, and EA represent a promising non-invasive therapeutic strategy for the management of drugs-caused neurotoxicity.

## MATERIALS AND METHODS

### Animals and treatments

Adult (~25 g, 8–10 weeks of age) male C57BL/6 mice were maintained on a 12-h reverse light-dark cycle with food and water available ad libitum. All mice were handled 5 min for 5 consecutive days preceding the experiment. Mice were assigned to receive a daily injection of METH (METH group, 3mg/kg in saline, i.p.) or Saline (SAL group, 0.2 mL, i.p.) for 10 consecutive days, then underwent 14-days withdrawal period at home cage. To determine the effect of electroacupuncture on METH-induced spatial memory deficits, EA treatments were performed at acupoints of GV29 and GV20 during 14-days METH withdrawal periods (METH+EA group); whereas sham EA (SEA) were performed at “placebo-point” (METH+SEA group) and the other operations were the same as the METH+EA group. All experiments were performed in accordance with Nanjing University of Chinese Medicine Guide for the Care and Use of Laboratory Animals, China. The study has been approved by the Nanjing University of Chinese Medicine Institutional Animal Care and Use Committee.

### Electroacupuncture treatments

24 h after the last METH injection, mice were fixed to a homemade plate with isoflurane (RWD, Shenzhen, China). For the EA treatments, two small acupuncture needles (0.25 × 13 mm, Hwato, China) were inserted gently in a depth of 2–3 mm to the acupoints at Baihui (GV 20, located in the midpoint between the auricular apices) and Yintang (GV29, located midway between the medial ends of the two eyebrows) ^17^, stimulated with dilatational waves of 2/100 Hz at an intensity of 1–2 mA (causing the slight vibration of muscles around acupoints) with peak voltage of 9 V by the EA device ^34^ (HANS-100A, Nanjing Jisheng Medical Technology, China) for 30 min, once a day for 14 consecutive days. For the SEA treatments: all procedures were same as the METH + EA group, the only difference was the acupoints located at “placebo-point” (below the costal region, 2 cm superior to the posterior superior iliac spine and ~3 cm lateral to the spine) ^35^.

### Morris water maze

The Morris water maze (MWM) test was performed to assess hippocampus-dependent spatial learning and memory. The experimental process included learning trials, escape latency test (T1) and probe test (T2). The platform was remained in the same position throughout the learning trials and T1, then removed from the pool during T2. Learning was measured by training mice to find a hidden platform (four trials per day, 90 s/trial) over 3 consecutive days of testing. If the mice did not find the platform within 90 s, they were guided to the platform. Once the mice reached the platform, they were allowed to stay on the platform for 15 s. Trials on each day of testing were averaged, and plotted as a time-course, with gradually decreasing escape latencies being used as our index of spatial learning behavior. T1 was conducted 24 h after the last day of training. Mice were placed in four quadrants in sequence, latency to enter the hidden platform were recorded. To assess spatial memory, T2 was performed in which the hidden platform was removed. Mice were placed in the pool from the diagonal quadrant of the target quadrant for one single 90-s probe test. Time spent exploring the target quadrant were quantified as measurements of spatial memory. All data were recorded and analyzed by TopScan H software.

### Y-maze

Y maze were completed and analyzed as previously described ^36^. Experiments were performed in a maze consisting of three transparent plastic arms, 40 × 10 × 25 cm each, set at a 120° angle relative to each other. Briefly, the Y-Maze comprised exposing mice to two arms (familiar arms, FA) of the maze for 5 min as a training trial. The third, blocked, arm of the maze was designated as the novel arm (NA), and was counterbalanced between mice. Following a 5-min delay, mice were returned to the maze but were free to explore all three arms for 5 min. TopScan H software was used for recording and analyzing the data. The time spent exploring the novel arm relative to the total time was quantified as our index of short-term spatial memory.

### Object location memory

Object location memory (OLM) tests were performed as previously described ^37^. Prior to the onset of training, mice were habituated to the training context during two 5-min sessions on two separate days. On the training trail, mice were placed in the experimental apparatus and allowed to explore two objects (a and b) for a total of three 5-min sessions with an intersession interval of 1–2 min. To avoid the presence of olfactory trails, the objects and apparatus were thoroughly cleaned with 70% ethanol between mice. 24 h after the training trial, object b was placed in a novel location (displaced object, DO), while object a was not moved (non-displaced object, NDO). Mice were placed in the experimental apparatus for 5 min, time spent exploring each object and total distance were recorded. Exploration of the objects was defined as pointing the nose to the object at a distance of <1 cm. Grooming near the object was not considered as exploration. Discrimination index was calculated as time spent exploring the DO relative to the total time spent exploring both objects (Discrimination index = time DO / (time DO + time NDO) x 100%). All data were recorded and analyzed by TopScan H software.

### Microdialysis

Mice were anesthetized with isoflurane after METH 14-days withdrawal. The micro-dialysis guide cannula (RWD, China) were implanted 1 mm above the dCA1 (AP −1.83mm, ML ±1.2mm, DV +1.6mm). Dental cement and a small metal peg were used to secure the guide cannula on the skull, then the duct cover were putted and tightened. After 3–4 d recovery period, microdialysis experiments were performed as previously described ^38^. Under isoflurane anesthesia, microdialysis probes (Probe length 4mm, membrane length 1mm and outer diameter 0.22 mm; EiCOM, AZ-4-01, USA) were inserted through the guide cannula in the dCA1 and perfused at a constant flow of 1.5 μL/min with artificial cerebrospinal fluid (aCSF, 140 mM NaCl, 3 mM KCl, 1.2-3.4 mM CaCl_2_, 1 mM MgCl_2_, 1.2 mM Na_2_HPO, 0.27 mM NaH_2_PO_4_, 7.2 mM glucose, pH = 7.4). Dialystates were collected for every 30 min and stored at −80 °C. At the end of the tests, histological verification of microdialysis probe location was performed.

### Liquid chromatography-mass spectrometry

The concentrations of Glu and Gln in the dCA1 were measured by liquid chromatography-mass spectrometry (LC-MS). Briefly, the dialysate samples were derivatized by adding 50 μL dansyl chloride solution (2 mg/mL, dissolved in acetone) and 50 μL sodium bicarbonate solution (pH = 9, 0.2 M NaHCO3) for 8 min in 60 *°C* water bath. After drying in a low temperature vacuum centrifugal enrichment system (Labconco), the residue was resuspended in 100 μL of acetonitrile. Samples were then centrifuged (29,700 g for 10 min) to remove additional precipitated proteins. Finally, sample (5 μL) was injected into the LC–MS system for further analysis. The LC-MS system was carried out using Thermo Vanquish F and TSQ Quantis system (Thermo Fisher Scientific). The internal standard (IS) was from the Institute for Food and Drug Control (Beijing, China). LC-MS was performed using Ulimate3000 ultra high-performance liquid chromatography system and Q Exactive four-stage pole orbital well high-resolution mass spectrometer (Thermo Fisher Scientific). The internal standard was from the Institute for Food and Drug Control (Beijing, China). Glu and Gln standards (Millipore Sigma, USA) was dissolved in methanol at concentrations of 1-200 ng/mL. Extracellular fluid Glu or Gln concentration in CA1 area = Glu or Gln concentration in dialysate / probe recovery.

### Immunofluorescence

The immunofluorescence was performed as previously described ^39^. Briefly, Mice were perfused with 0.9% Saline followed by 4% paraformaldehyde in PBS buffer (pH 7.4). The brains were removed, post-fixed in 4% paraformaldehyde for 2 h, and then transferred to 30% sucrose overnight. Frozen coronal sections (30 μm) were cut on cryostats. Coronal brain slices containing dorsal hippocampus were incubated with primary antibody overnight at 4°C. The following primary antibodies were used in these experiments: rat anti-C3 (1:200; abcam, UK), mouse anti-S100A10 (1:200; Thermo Fisher Scientific, USA), rabbit anti-GFAP (1:300; Boster, China), rat anti-GFAP(1:500, Thermo Fisher Scientific, USA), rabbit anti-GLT-1 (1:300; Cell Signaling Technology, USA), rabbit anti-GLAST (1:300; Thermo Fisher Scientific, USA), mouse anti-GS (1:500; abcam, UK), rabbit anti-c-Fos (1:2000; Cell Signaling Technology, USA), rabbit anti-SYP1 (Synaptotagmin-1, 1;500; Signalway Antibody, USA), mouse anti-VGLUT-1 (1;500; 1:200; abcam, UK), rabbit anti-NeuN (1:100; MERCK, Germany). Then the sections were rinsed in PBS, and subsequently incubated with secondary antibody for 1 h at room temperature. The following secondary antibodies were used here: Alexa Fluor 555-labeled donkey anti rabbit secondary antibody (1:500, Thermo Fisher Scientific, USA), Alexa Fluor 555-labeled donkey anti rat secondary antibody (1:500, Thermo Fisher Scientific, USA) Alexa Fluor 488-labeled donkey anti-mouse secondary antibody (1:500; Thermo Fisher Scientific, USA). Negative controls were performed by replacing primary antibodies with PBS buffer. The images were captured by Leica DM6 B upright digital research microscope (Leica Microsystems CMS GmbH, Germany).

### Western blot

Total protein was extracted from the dCA1. Protein was electrophoretically transferred onto PVDF membranes. 20 ng per microliter protein was loaded in the gel ^39^. The blots were incubated overnight at 4°C with the primary antibody [rat anti-C3 (1:50; abam, UK), rabbit anti-GFAP (1:1000; Boster, China), rabbit anti-GLT-1 (1:1000; Thermo Fisher Scientific, USA), rabbit anti-GLAST (1:1000; abcam, UK), mouse anti-GS (1:1000; abcam, UK), mouse anti-S100A10 (1:1000; Thermo Fisher Scientific, USA). The blots were then washed and incubated with HRP-conjugated secondary antibody (Boster, China) for 1 h at room temperature and observed using an ECL kit (Cwbio, Beijing, China). Proteins were normalized to β-actin (1:20000; proteintech, USA). The blots were washed with stripping buffer (Beyotime Institute of Biotechnology) to allow reprobing with other antibodies. Values for protein levels were calculated with Image J.

### Primary astrocyte culture and treatment

Primary cortical astrocytes were prepared using 1 - day old C57BL/6 mice. The mice were decapitated and the cerebral cortices were removed carefully. Tissue pieces were transferred to 0.25% trypsin-EDTA solution (Thermo Fisher Scientific) followed by 10 min of gentle agitation. Tissues were maintained in DMEM and nutrient mixture F12 (DMEM/F12) containing 10% FBS and dissociated into single cells. The cultures were incubated at 37°C in a humidified 5% CO2, 95% air atmosphere. The medium was changed every 3 days, until reaching confluence, which usually occurred after 7–10 days. Before changing the medium. The flasks were shaken at 180 rpm for 30 min on an orbital shaker to remove microglia and discarded the supernatant containing microglia. Afterwards, the purification of astrocytes was carried out. Cells were treated with serum-free medium 0.25% trypsin at 37°C, then added the same amount of medium to stop trypsinization.

Mixed the liquid in the tube and centrifuged at 1000 r/min for 5 min. Cells were then resuspended in 10 mL of medium and cultured in a well-prepacked flask for 1 d. Finally, the flask was shaken at 200 rpm for 2h at 37 ° C. After two purification processes, purified astrocyte cultures were obtained, which were constituted by more than 95% of GFAP - positive cells. A1 reactive astrocytes were generated in vitro by growing purified astrocytes for 6 days and then treated with Il-1α (3 ng/mL, STEMCELL, 78115), TNFα (30 ng/mL, Cell Signaling Technology, 8902SF), and C1q (400 ng/mL, abcam, ab96363) for 24 h ^12^.

### Primary astrocyte staining

Astrocytes were seeded at 4.0 × 10^5^ on 3.0 mm^2^ coverslips for 24 h. The cells were fixed with 4% icecold paraformaldehyde for 10 min at 4°C. The cells were then air dried, followed by blocking and permeabilization with 1% BSA in PBS with 0.1% Triton for 1 h. After that, astrocyte (A1) was stained with C3 antibody (1:200) and GFAP antibody (1:300) 24 h in a humidified chamber at 4°C. After three washes with 0.1% Triton in PBS, goat anti - rabbit IgG H&L (Alexa Fluor 555, 1:500) and goat anti - rat IgG H&L (Alexa Fluor 488, 1:500) was used for 1 h. After DAPI staining for 10 min, the coverslips were transferred onto glass slides. Images were obtained using a digital microscope camera system (Leica Microsystems, Buffalo Grove, IL, USA).

### Real-time reverse transcription polymerase chain reaction (RT-PCR)

Total RNA was extracted using a FastPure Cell/Tissue Total RNA Isolation Kit (Cat.RC101-01, Vazyme, China), according to the manufacturer’s instructions. Reverse transcriptions were performed using a miRNA 1st Strand cDNA Synthesis Kit (MR101-01/02, Vazyme, China). The C3 and GAPDH primers were as follows: C3 (forward, GAGCGAAGAGACCATCGTACT; reverse, TCTTTAGGAAGTCTTGCACAGTG); GAPDH (forward, TCATCCCAGCTGAACG; reverse, TCATACTTGGCAGGTTTCTCC;). Target gene expression was normalized to GAPDH expression, GAPDH was used as the internal control. The relative expression level was calculated by the comparative CT method (2-ΔΔCt).

### Glutamate uptake assay

Glutamate clearance capacity was measured using Amplex™ Red Glutamic Acid/Glutamate Oxidase Assay Kit (Thermo Fisher Scientific, USA) following manufacturer’s protocol. The culture medium was replaced with 2 mL serum-free HBSS containing 100 μM glutamate (Sigma, USA) after A1 astrocytes induction ^40^. Cells were returned to the incubator for 1 h at 37 °C, then 50 μL of supernatant was transferred to a tube; 2 μl of supernatant was added to 96-well plate and determined the remaining glutamate in the medium. The absorbance at 530 nM was obtained using Multi-function enzyme marker workstation (Molecular Devices, USA). The concentration in the supernatant was calculated using the standard curve, and calculated the changes of glutamate concentration.

### Patch-clamp electrophysiology

Mice were deeply anesthetized with isoflurane (RWD, Shenzhen, China) before receiving a perfusion of 10 mL ice-cold cutting solution containing the following: 135 mM NMDG, 1 mM KCl, 1.2 mM KH_2_P0_4_, 1.5 mM MgCl_2_·6H_2_O, 0.5 mM CaCl_2_·2H_2_O, 20 mM Choline Bicarbonate, 10 mM D-(+)- Glucose, pH adjusted to 7.4 with HCl, and saturated with 95% O_2_ / 5% CO_2_. Coronal slices of 300 μm thickness were cut containing the dCA1 using a vibratome (Leica VT1000 S, Germany) in ice-cold cutting solution. Slice were allowed to recover in oxygenated artificial cerebrospinal fluid (aCSF) [containing (in 119 mM NaCl, 2.5 mM KCl, 1 mM NaH_2_PO_4_O_2_H_2_O, 26.2 mM NaHCO, 1.3 mM MgCl_2_, 2.5 mM CaCl_2_, 11 mM D-Glucose, pH7.4) first in 37°C for 30 min and then at room temperature for a total of 1 h. For recordings, one slice was transferred from the holding chamber to a recording chamber, where it was continuously perfused with oxygenated aCSF. A cesium internal solution (130 mM CsMeSO4, 10 mM NaCl, 10 mM EGTA, 4 mM MgATP, 0.3 mM Na3GTP, 10 mM HEPES, pH7.4) was used for paired-pulse ratio (PPR) and miniature EPSCs (mEPSCs) recording. For PPR measurements, picrotoxin (100 μM) was added to the recording aCSF and a stimulating electrode was placed at the Schaffer fibers in the dCA1. For PPR, every 15 s, a paired-pulse stimulation was delivered with a 20, 50 or 100 ms inter-stimulus intervals (ISI). Stimulation duration was 0.1 ms and current was approximately 0.02-0.08 mA. Evoked EPSCs were recorded in the dCA1 neurons at a holding potential of −70 mV. 5 repetitions of each ISI were recorded per cell after stable evoked EPSCs was achieved. PPR was calculated as the ratio between the peak amplitudes of the second and first EPSC. mEPSCs were recorded in the presence of TTX (1 μM), d-AP5 (50 μM) and Picrotoxin (100 μM) in the recording aCSF solution. All recordings swere filtered (4 kHz low-pass filter) and sampled (10 kHz) for online and later offline analysis with Multiclamp 700B amplifier (Molecular Devices, Sunnyvale, CA, USA) and Clampfit v.10.5 software (Molecular Devices).

### Fiber photometry in Y maze

A mixture of viruses (AAV2/9-CaMKⅡ-VRE-WPRE-pA and AAV2/9-EF1a-DIO-axon-Gcamp6s-WPRE-pA, 1:3, Brain VTA Technology Co, China) were injected into the dCA3 (AP −1.83mm, ML ± 2.75mm, DV +1.6mm). Afterwards an optic fiber was implanted in the dCA1 (AP −1.83mm, ML ±1.2mm, DV +1.6mm), and the catheter was fixed on the skull with dental cement and a small metal screw. After 7–8 d recovery period, mice were assigned to receive a daily injection of METH or Saline for 10 consecutive days, then underwent 14-days withdrawal period at home cage. On day 30, the realtime presynaptic (dCA3) axonal calcium signals were recorded in the dCA1during Y maze test with an optical fiber recording system (Thinker Tech Nanjing Biotech Limited Co., Ltd). A 488 nm laser (0.01–0.02 mW) was delivered using an optical fiber recording system, and fluorescent signals were recorded. Fluorescence heat maps were calculated by binning the GCaMP fluorescence as a function of mouse position. For data analysis, we set the time point at which the mouse entered the novel arm (NA) as “0”. Matlab was used to analyze the change in fluorescence intensity before and after “0”.

### Statistical analysis

All data are expressed as means ± SEM. Statistical analysis was carried out using Prism v.8.0 (GraphPad Software, La Jolla, CA, USA). Figure 3 A3, Figure 1 D1 and Figure 4 C were analyzed by 2-way ANOVA followed by Bonferroni’s post hoc test, others were analyzed by 2-tailed Student’s t test. Statistical significance was set at P < 0.05.

## AUTHOR CONTRIBUTIONS

PBS, ZSL, XX and JXN performed the majority of the experiments, they are all co–first authors. Their order in the author list was based on the first key experiments, which were performed and analyzed by PBS. ZSL performed primary astrocyte culture and treatment. XX and JXN performed patch-clamp electrophysiology and fiber photometry. DKL and QLC helped with behavioral studies in mice. YHZ collected and analyzed animal data. YG provided technical training and edit the manuscript. XWG and FFG developed the overall concept, supervised the experiments, analyzed data, and wrote the manuscript.

## CONFLICTS OF INTEREST

The authors declare no competing interests.

## ACKNOWLEDGMENTS

This work is supported by National Natural Science Foundation of China (No. 81571303 and 81901353), and the Priority Academic Program Development of Jiangsu Higher Education Institutions (Integration of Chinese and Western Medicine).

## Notes

### Competing Interest Statement

The authors have declared no competing interest.

## References

1. Szumlinski, KK, Lominac, KD, Campbell, RR, Cohen, M, Fultz, EK, Brown, CN, et al. (2017). Methamphetamine Addiction Vulnerability: The Glutamate, the Bad, and the Ugly. Biol Psychiatry 81: 959–970.

2. Cox, BM, Cope, ZA, Parsegian, A, Floresco, SB, Aston-Jones, G, and See, RE (2016). Chronic methamphetamine self-administration alters cognitive flexibility in male rats. Psychopharmacology (Berl) 233: 2319–2327.

3. McKetin, R, Lubman, DI, Baker, AL, Dawe, S, and Ali, RL (2013). Dose-related psychotic symptoms in chronic methamphetamine users: evidence from a prospective longitudinal study. JAMA Psychiatry 70: 319–324.

4. Mishra, D, Pena-Bravo, JI, Leong, KC, Lavin, A, and Reichel, CM (2017). Methamphetamine self-administration modulates glutamate neurophysiology. Brain Struct Funct 222: 2031–2039.

5. Lisman, J, Buzsaki, G, Eichenbaum, H, Nadel, L, Ranganath, C, and Redish, AD (2017). Viewpoints: how the hippocampus contributes to memory, navigation and cognition. Nat Neurosci 20: 1434–1447.

6. Yu, JY, Fang, P, Wang, C, Wang, XX, Li, K, Gong, Q, et al. (2018). Dorsal CA1 interneurons contribute to acute stress-induced spatial memory deficits. Neuropharmacology 135: 474–486.

7. Golsorkhdan, SA, Boroujeni, ME, Aliaghaei, A, Abdollahifar, MA, Ramezanpour, A, Nejatbakhsh, R, et al. (2020). Methamphetamine administration impairs behavior, memory and underlying signaling pathways in the hippocampus. Behav Brain Res 379: 112300.

8. Alshehri, FS, Althobaiti, YS, and Sari, Y (2017). Effects of Administered Ethanol and Methamphetamine on Glial Glutamate Transporters in Rat Striatum and Hippocampus. J Mol Neurosci 61: 343–350.

9. Siemsen, BM, Reichel, CM, Leong, KC, Garcia-Keller, C, Gipson, CD, Spencer, S, et al. (2019). Effects of Methamphetamine Self-Administration and Extinction on Astrocyte Structure and Function in the Nucleus Accumbens Core. Neuroscience 406: 528–541.

10. Bernardinelli, Y, Randall, J, Janett, E, Nikonenko, I, Konig, S, Jones, EV, et al. (2014). Activity-dependent structural plasticity of perisynaptic astrocytic domains promotes excitatory synapse stability. Curr Biol 24: 1679–1688.

11. Liddelow, SA, and Barres, BA (2017). Reactive Astrocytes: Production, Function, and Therapeutic Potential. Immunity 46: 957–967.

12. Liddelow, SA, Guttenplan, KA, Clarke, LE, Bennett, FC, Bohlen, CJ, Schirmer, L, et al. (2017). Neurotoxic reactive astrocytes are induced by activated microglia. Nature 541: 481–487.

13. Coulter, DA, and Eid, T (2012). Astrocytic regulation of glutamate homeostasis in epilepsy. Glia 60: 1215–1226.

14. Rose, CR, Felix, L, Zeug, A, Dietrich, D, Reiner, A, and Henneberger, C (2017). Astroglial Glutamate Signaling and Uptake in the Hippocampus. Front Mol Neurosci 10: 451.

15. Jang, JH, Kim, YK, Jung, WM, Kim, HK, Song, EM, Kim, HY, et al. (2019). Acupuncture Improves Comorbid Cognitive Impairments Induced by Neuropathic Pain in Mice. Front Neurosci 13: 995.

16. Li, G, Zeng, L, Cheng, H, Han, J, Zhang, X, and Xie, H (2019). Acupuncture Administration Improves Cognitive Functions and Alleviates Inflammation and Nuclear Damage by Regulating Phosphatidylinositol 3 Kinase (PI3K)/Phosphoinositol-Dependent Kinase 1 (PDK1)/Novel Protein Kinase C (nPKC)/Rac 1 Signaling Pathway in Senescence-Accelerated Prone 8 (SAM-P8) Mice. Med Sci Monit 25: 4082–4093.

17. Zheng, Y, He, J, Guo, L, Yao, L, Zheng, X, Yang, Z, et al. (2019). Transcriptome Analysis on Maternal Separation Rats With Depression-Related Manifestations Ameliorated by Electroacupuncture. Front Neurosci 13: 314.

18. Zhang, M, Xv, GH, Wang, WX, Meng, DJ, and Ji, Y (2017). Electroacupuncture improves cognitive deficits and activates PPAR-gamma in a rat model of Alzheimer’s disease. Acupunct Med 35: 44–51.

19. Lin, L, Yu, L, Xiang, H, Hu, X, Yuan, X, Zhu, H, et al. (2019). Effects of Acupuncture on Behavioral Stereotypies and Brain Dopamine System in Mice as a Model of Tourette Syndrome. Front Behav Neurosci 13: 239.

20. Escartin, C, Guillemaud, O, and Carrillo-de Sauvage, MA (2019). Questions and (some) answers on reactive astrocytes. Glia 67: 2221–2247.

21. Lominac, KD, Sacramento, AD, Szumlinski, KK, and Kippin, TE (2012). Distinct neurochemical adaptations within the nucleus accumbens produced by a history of self-administered vs non-contingently administered intravenous methamphetamine. Neuropsychopharmacology 37: 707–722.

22. Yang, W, Yang, R, Luo, J, He, L, Liu, J, and Zhang, J (2018). Increased Absolute Glutamate Concentrations and Glutamate-to-Creatine Ratios in Patients With Methamphetamine Use Disorders. Front Psychiatry 9: 368.

23. Halpin, LE, Northrop, NA, and Yamamoto, BK (2014). Ammonia mediates methamphetamine-induced increases in glutamate and excitotoxicity. Neuropsychopharmacology 39: 1031–1038.

24. Pena-Bravo, JI, Penrod, R, Reichel, CM, and Lavin, A (2019). Methamphetamine Self-Administration Elicits Sex-Related Changes in Postsynaptic Glutamate Transmission in the Prefrontal Cortex. eNeuro 6.

25. Pirttimaki, TM, Sims, RE, Saunders, G, Antonio, SA, Codadu, NK, and Parri, HR (2017). Astrocyte-Mediated Neuronal Synchronization Properties Revealed by False Gliotransmitter Release. J Neurosci 37: 9859–9870.

26. Scofield, MD (2018). Exploring the Role of Astroglial Glutamate Release and Association With Synapses in Neuronal Function and Behavior. Biol Psychiatry 84: 778–786.

27. Kruyer, A, Scofield, MD, Wood, D, Reissner, KJ, and Kalivas, PW (2019). Heroin Cue-Evoked Astrocytic Structural Plasticity at Nucleus Accumbens Synapses Inhibits Heroin Seeking. Biol Psychiatry 86: 811–819.

28. Zhou, M, and Kimelberg, HK (2001). Freshly isolated hippocampal CA1 astrocytes comprise two populations differing in glutamate transporter and AMPA receptor expression. J Neurosci 21: 7901–7908.

29. Zhou, Y, Wang, X, Tzingounis, AV, Danbolt, NC, and Larsson, HP (2014). EAAT2 (GLT-1; slc1a2) glutamate transporters reconstituted in liposomes argues against heteroexchange being substantially faster than net uptake. J Neurosci 34: 13472–13485.

30. Scofield, MD, Heinsbroek, JA, Gipson, CD, Kupchik, YM, Spencer, S, Smith, AC, et al. (2016). The Nucleus Accumbens: Mechanisms of Addiction across Drug Classes Reflect the Importance of Glutamate Homeostasis. Pharmacol Rev 68: 816–871.

31. Litscher, G (2019). Ear Acupuncture according to the NADA (National Acupuncture Detoxification Association). Medicines (Basel) 6.

32. Zhang, Y, Cai, XH, Zhang, RJ, Hou, XR, Song, XG, Wu, SB, et al. (2016). Acupuncture regulates the unfolded protein response and inhibits apoptosis in a rat model of heroin relapse. Acupunct Med 34: 441–448.

33. Cao, J, Tang, Y, Li, Y, Gao, K, Shi, X, and Li, Z (2017). Behavioral Changes and Hippocampus Glucose Metabolism in APP/PS1 Transgenic Mice via Electro-acupuncture at Governor Vessel Acupoints. Front Aging Neurosci 9: 5.

34. Li, XY, Xu, L, Liu, CL, Huang, LS, and Zhu, XY (2016). [Electroacupuncture Intervention Inhibits the Decline of Learning-memory Ability and Overex-pression of Cleaved Caspase-3 and Bax in Hippocampus Induced by Isoflurane in APPswe/PS 1]. Zhen Ci Yan Jiu 41: 24–30.

35. Liu, W, Zhuo, P, Li, L, Jin, H, Lin, B, Zhang, Y, et al. (2017). Activation of brain glucose metabolism ameliorating cognitive impairment in APP/PS1 transgenic mice by electroacupuncture. Free Radic Biol Med 112: 174–190.

36. Notaras, M, Allen, M, Longo, F, Volk, N, Toth, M, Li Jeon, N, et al. (2019). UPF2 leads to degradation of dendritically targeted mRNAs to regulate synaptic plasticity and cognitive function. Mol Psychiatry.

37. Hall, JH, Wiseman, FK, Fisher, EM, Tybulewicz, VL, Harwood, JL, and Good, MA (2016). Tc1 mouse model of trisomy-21 dissociates properties of short-and long-term recognition memory. Neurobiol Learn Mem 130: 118–128.

38. Kinoshita, H, Nishitani, N, Nagai, Y, Andoh, C, Asaoka, N, Kawai, H, et al. (2018). Ketamine-Induced Prefrontal Serotonin Release Is Mediated by Cholinergic Neurons in the Pedunculopontine Tegmental Nucleus. Int J Neuropsychopharmacol 21: 305–310.

39. Shi, P, Nie, J, Liu, H, Li, Y, Lu, X, Shen, X, et al. (2019). Adolescent cocaine exposure enhances the GABAergic transmission in the prelimbic cortex of adult mice. FASEB J 33: 8614–8622.

40. Cho, IK, Yang, B, Forest, C, Qian, L, and Chan, AWS (2019). Amelioration of Huntington’s disease phenotype in astrocytes derived from iPSC-derived neural progenitor cells of Huntington’s disease monkeys. PLoS One 14: e0214156.

