## Supplementa Figures and legends for "Electro-acupuncture Alleviates METH Withdrawal-induced Spatial Memory Deficits by Restoring Astrocyte-drived Glutamate Uptake in dCA1"

**Supplemental Figure 1.** EA treatments did not affect METH-impaired object location memory (OLM) in mice. A, Experimental design of the OLM task. B, Discrimination index of OLM test. Discrimination index was calculated as time spent exploring the displaced object (DO, b) relative to the total time spent exploring the two objects [DO + non-displaced object (NDO, a), i.e., Discrimination index = time DO / (time DO + time NDO) x 100%]. C, Total distance traveled during the OLM test. Data are Mean ± S.E.M.

**Supplemental Figure 2.** METH and EA treatments did not alter glutamine (Gln) concertation in the dCA1. A, Gln levels in the dCA1of SAL and METH mice. B, Gln levels in the dCA1 of M+SEA and M+EA mice. Data are Mean ± S.E.M.

**Supplemental Figure 3.** METH and EA treatments did not change A2-like astrocytes in the dCA1. A, S100A10 staining in the dCA1 during METH withdrawal. Left, immunohistochemistry micrographs for S100A10 (green), GFAP (red) and DAPI (blue); Right, the ratio of colocalization of S100A10 and GFAP. B, Western blots (top) and quantification (bottom) of S100A10 protein levels in the dCA1. Data are Mean ± S.E.M.

**Supplemental Figure 4.** GS was highly enriched in astrocytes but not in neurons. A, Immunohistochemistry images for GS (green), NeuN (red) and DAPI (blue). B, Quantification of colocalization of GS with neuron (NeuN) and astrocyte activation marker (GFAP). Data are Mean ± S.E.M.

### Supplemental Figure 1

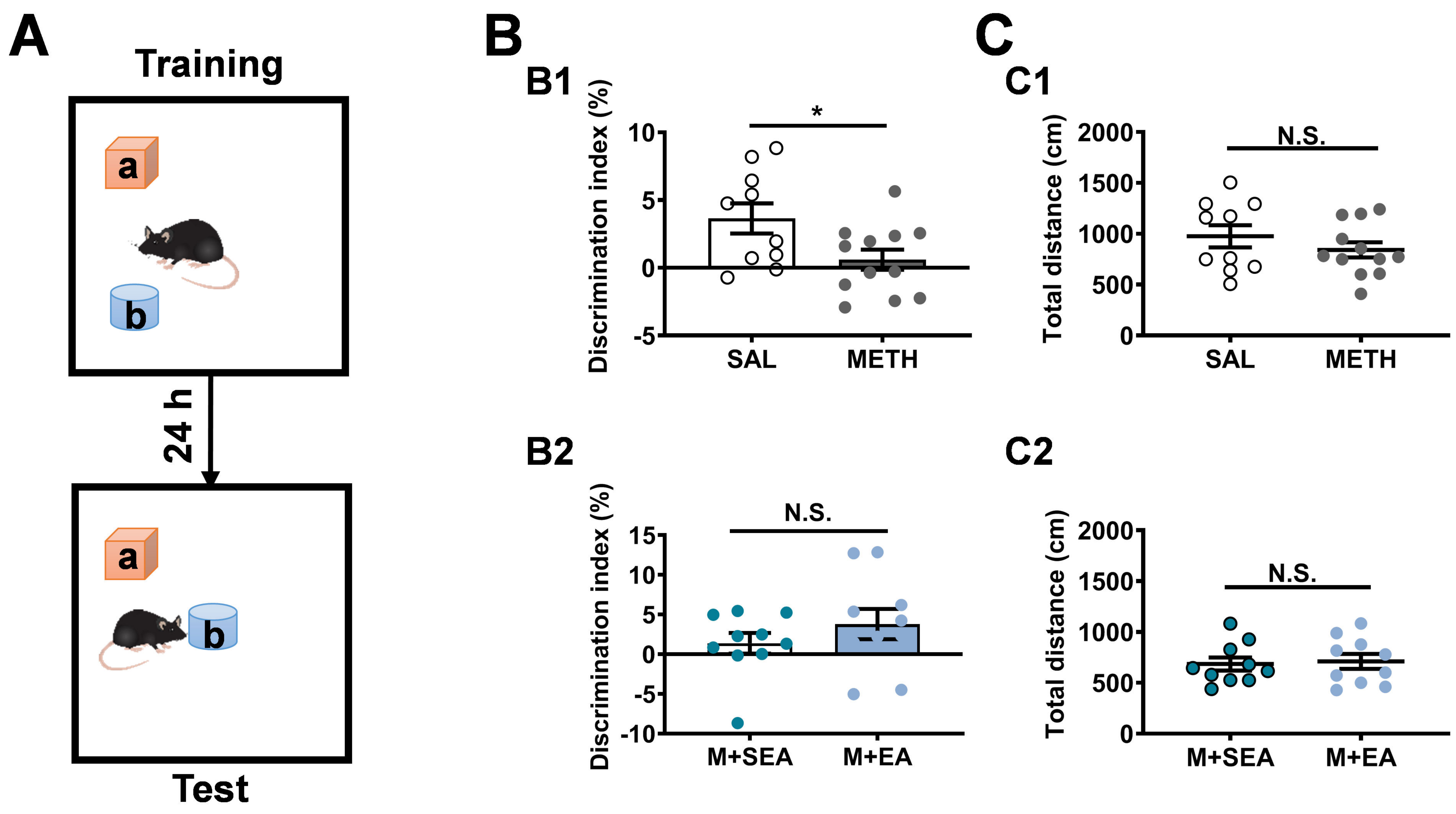

### Supplemental Figure 2

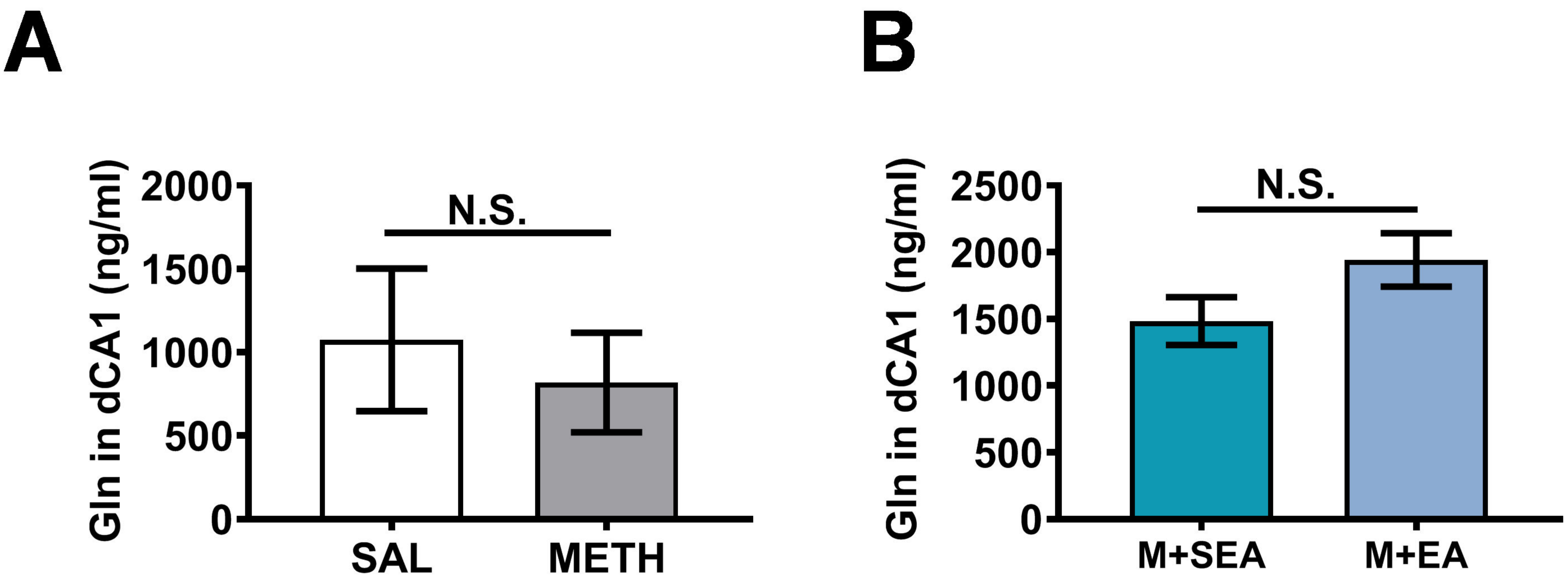

### Supplemental Figure 3

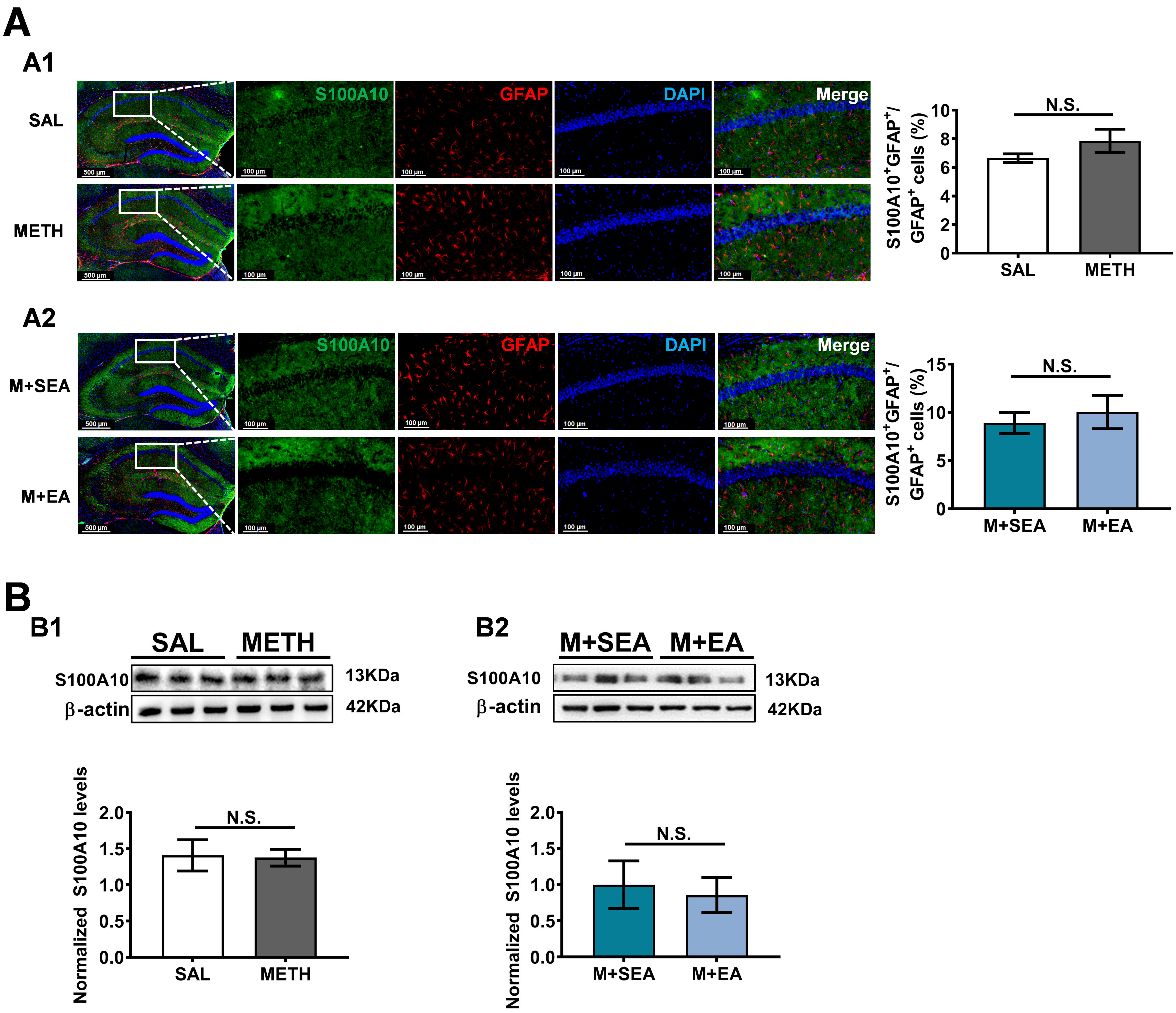

### Supplemental Figure 4

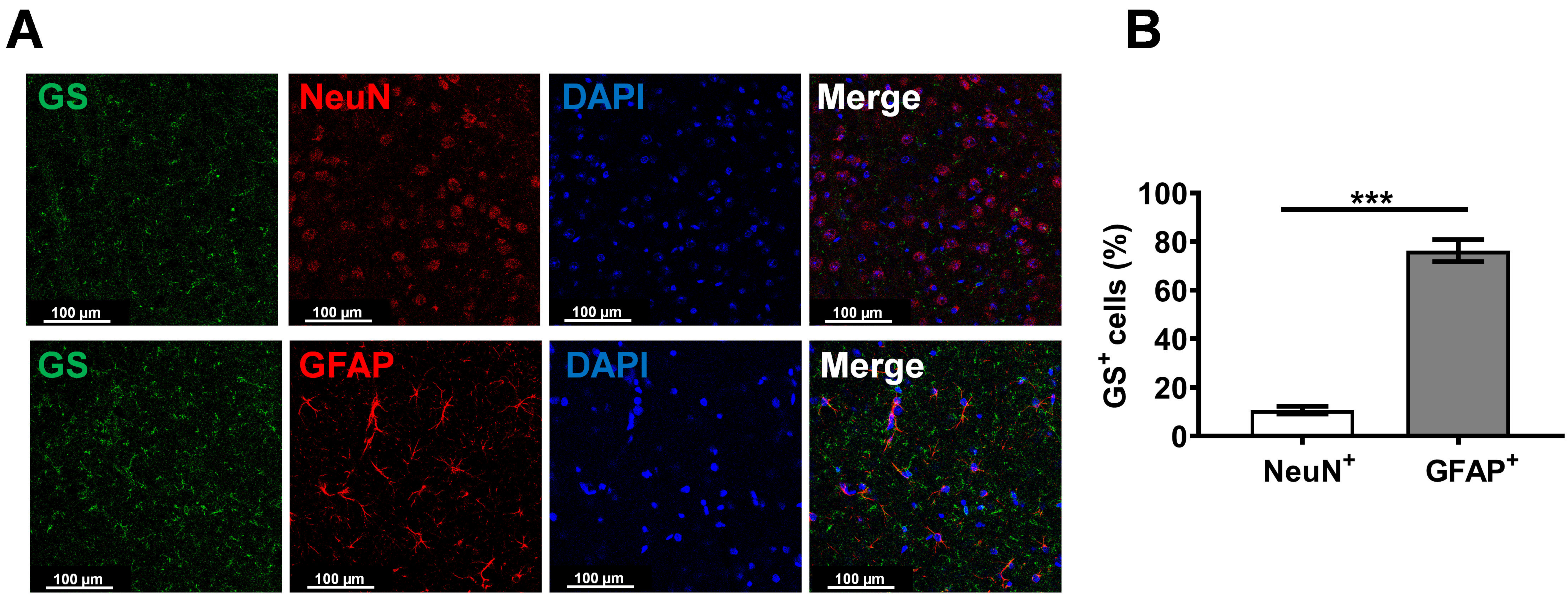
